# mosGraphGPT: a foundation model for multi-omic signaling graphs using generative AI

**DOI:** 10.1101/2024.08.01.606222

**Authors:** Heming Zhang, Di Huang, Emily Chen, Dekang Cao, Tim Xu, Ben Dizdar, Guangfu Li, Yixin Chen, Philip Payne, Michael Province, Fuhai Li

**Author notes:** Co-first authors (Heming Zhang, Di Huang).

## Abstract

Generative pretrained models represent a significant advancement in natural language processing and computer vision, which can generate coherent and contextually relevant content based on the pre-training on large general datasets and fine-tune for specific tasks. Building foundation models using large scale omic data is promising to decode and understand the complex signaling language patterns within cells. Different from existing foundation models of omic data, we build a foundation model, *mosGraphGPT*, for multi-omic signaling (mos) graphs, in which the multi-omic data was integrated and interpreted using a multi-level signaling graph. The model was pretrained using multi-omic data of cancers in The Cancer Genome Atlas (TCGA), and fine-turned for multi-omic data of Alzheimer’s Disease (AD). The experimental evaluation results showed that the model can not only improve the disease classification accuracy, but also is interpretable by uncovering disease targets and signaling interactions. And the model code are uploaded via GitHub with link: https://github.com/mosGraph/mosGraphGPT

## 1. Introduction

Generative pretrained models have significantly advanced fields such as natural language processing and computer vision^1^. These models are initially trained on extensive datasets and later fine-tuned for specific tasks, allowing them to produce coherent and contextually relevant content^2^. In bioinformatics, the need for foundational models arises due to the complexity and volume of biological data^3^. Traditional models, like SVM or Autoencoders, often struggle with the variability in gene expression and the diverse conditions of cell types^4^. Foundation models overcome these challenges by learning generalized representations from large-scale datasets, capturing complex gene-gene and gene-cell interactions that simpler models cannot^5^. Additionally, foundation models benefit from extensive pretraining on massive datasets, efficiently extracting key features and outperforming traditional models that typically cannot generalize across different contexts without requiring extensively labeled datasets for specific tasks^6^.

Advancements in sequencing technologies have led to the generation of multi-omic data^7,8^, which is essential for understanding the genetic diversity and complex signaling pathways at various levels within diseases, including cancer and Alzheimer’s Disease (AD). The multi-omic datasets of cancer and AD are publicly available. However, the integrative multi-omic data analysis remains an open problem to identify the essential (sparse a few) signaling targets and signaling pathways from thousands of targets densely interacting with each other, interpreting the molecular mechanisms and novel therapeutic targets. AD is commonly defined using criteria such as the CERAD (Consortium to Establish a Registry for Alzheimer’s Disease) score^9^, which evaluates the density of neurotic plaques to classify the severity of the disease. Many reports of omics data and analyses of AD have been published^10–25^. However, the pathogenesis of AD remains unclear and there is a lack of effective prevention and curable treatment medications.

Compared to single-omic data analysis, integrating multi-omic datasets offers a comprehensive perspective on intricate and multi-layered biological processes. This integration enhances statistical power, enabling the identification of molecular mechanisms that involve crucial molecular targets and signaling pathways^26^. Multi-omics data integration can improve the prediction and understanding of these conditions by revealing the genetic, transcriptomic, proteomic, and metabolomic alterations associated with disease progression and severity^27^.

Proteins within cells function as part of systematic networks and modules, regulating complex biological processes and dysfunctional signaling pathways in diseases such as cancer^28^. Several signaling pathways, such as those documented in KEGG^29^, WikiPathways^30,31^, and protein-protein interaction (PPI) databases like BioGRID^32,33^ and STRING^34,35^, are publicly accessible. Graph neural network (GNN)-based models can effectively represent the flow and interactions within these signaling networks. The latent state of individual proteins is influenced by their multi-omics data features and the interacting proteins (neighbors) within the signaling network. Importantly, attention mechanisms can be employed to identify crucial targets and subsequent signaling pathways. Several models have been developed to integrate multi-omics data for a deeper understanding of complex diseases such as AD and Non-AD. Among them, M3NetFlow^14^ is a sophisticated model designed to incorporate multi-omics data into a graph-based framework. M3NetFlow leverages multi-hop information within each subgraph and employs global bi-directional message propagation to facilitate communication between genes and proteins. This approach enhances the inference process, allowing for a more nuanced understanding of the underlying biological processes^36^. Another noteworthy model is mosGraphGen (multi-omics signaling graph generator)^12^, which generates multi-omics signaling graphs for individual samples. This tool maps multi-omics data onto a biologically meaningful multi-level signaling network, enabling integrative and interpretable multi-omics data analysis using GNN models. By constructing these detailed signaling graphs, mosGraphGen provides valuable insights into the multi-layered interactions within the biological systems of individual samples.

Building upon the strengths of these models, we propose a novel approach that further enhances the integration and analysis of multi-omics data. The graph foundation model aims to address the limitations of existing models by incorporating advanced generative pre-trained models and graph neural networks. By leveraging the extensive pretraining capabilities of foundation models, our approach can capture complex gene-gene and gene-cell interactions with higher accuracy and contextual relevance. In this study, we pre-trained the model using TCGA cancer multi-omic data^37^, and fine-turned the model using multi-omic data of AD to identify the key targets and signaling pathways of AD.

## 2. Methodology and Materials

### 2.1. Datasets

#### Multi-omics datasets of Alzheimer’s Disease

The multi-omics data can be obtained from publicly available datasets, UCSC Xena and ROSMAP datasets (see **Tables 1-2**). After downloading the multi-omics data (including epigenomics, genomics, transcriptomics, proteomics, etc.) from these sources, the datasets will be converted into 2-dimensional data frames. These data frames will have columns for sample IDs, sample names, etc., and rows for probes, gene symbols, gene IDs, etc. To integrate multi-omics data with clinical data, identical samples across the datasets must be identified. Similarly, the process must involve converting rows (probes, gene symbols, gene IDs, etc.) into an identical standard: gene-level data by either aggregating the same measurements for each gene or eliminating duplicates due to gene synonyms. Genes are then aligned according to a reference genome, ensuring that the final annotation for each gene in the multi-omics data is accurate. Finally, standardizing gene counts across multi-omics datasets and addressing missing values by imputing with zeros or negative one values where necessary. After aligning all the columns to standard sample IDs and all the rows to standard gene IDs and unifying identical number of samples and genes, the data were prepared for integration into Graph Neural Network (GNN) models. The epigenomics, genomics, transcriptomics, and proteomics data will be utilized as features of protein nodes within the GNN models.

**Table 1.**
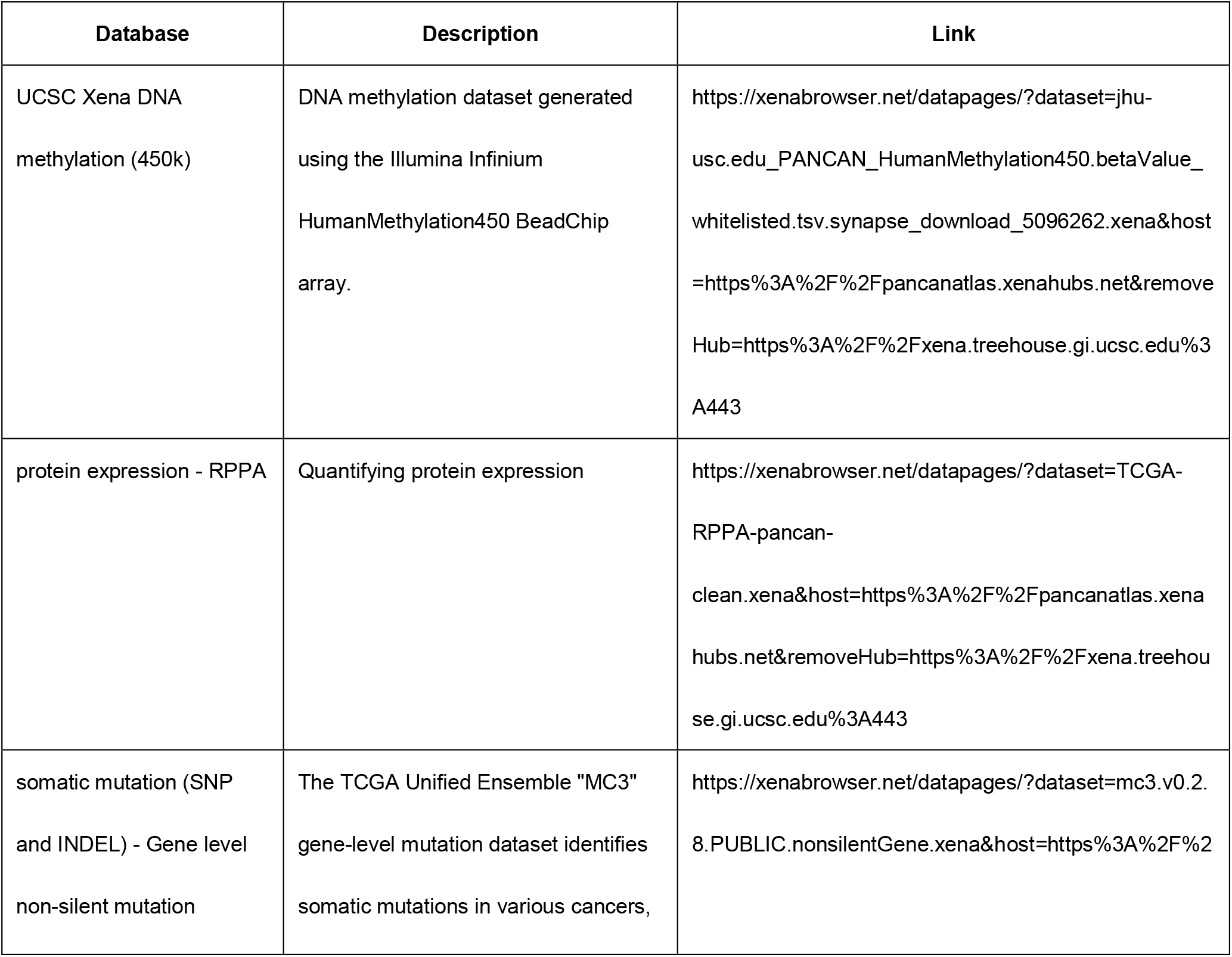
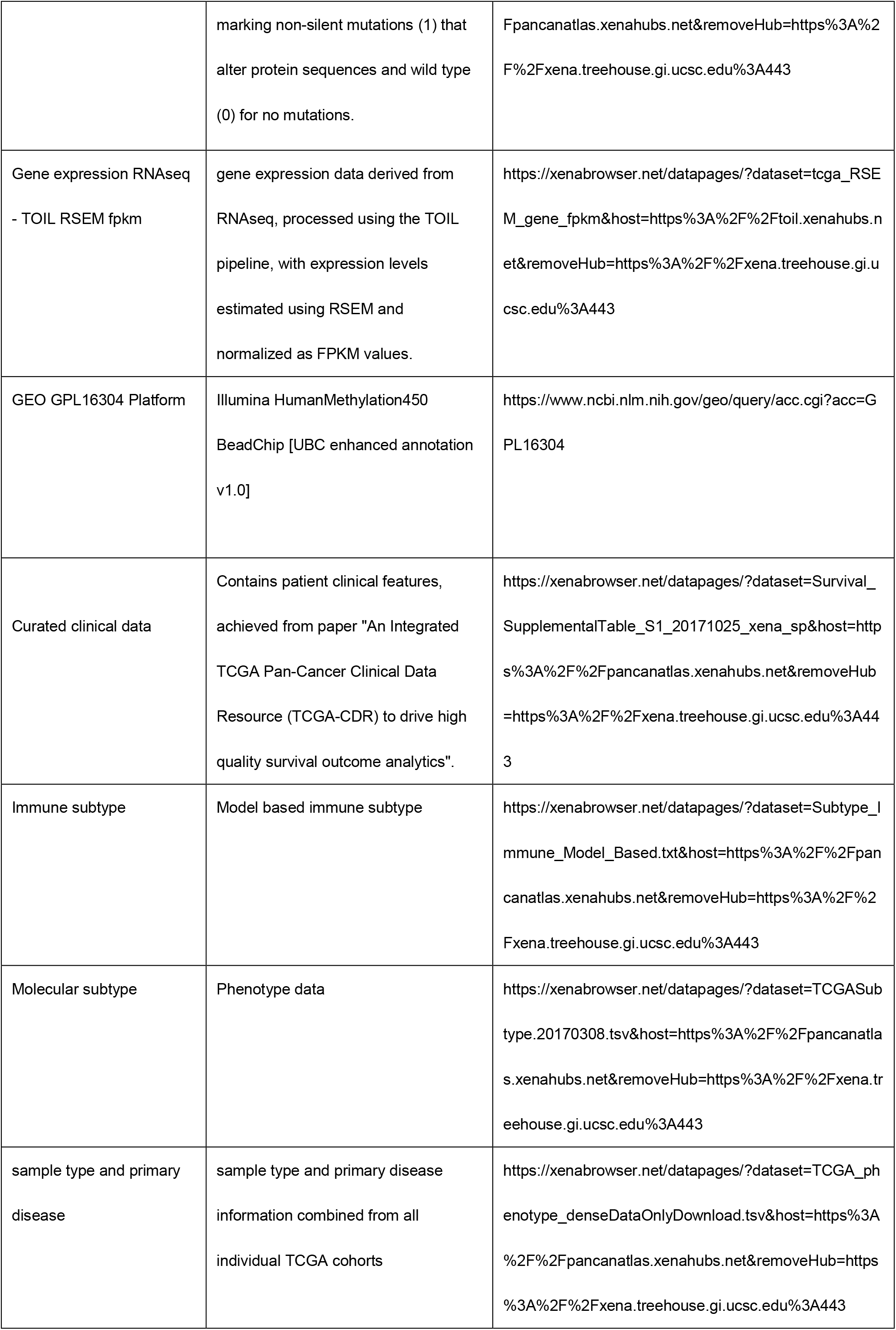
UCSC Database resources.

**Table 2.** ROSMAP Database resources.

| Database | Description | Link |
| --- | --- | --- |
| ROSMAP_arrayMethylation_imputed | Methylation data was generated on prefrontal cortex samples collected from 708 individuals using the Illumina HumanMethylation450 BeadChip | https://www.synapse.org/#!Synapse:syn3168763 |
| C2.median_polish_corrected_log2(Proteomics) | Data generated from isobaric TMT peptide labeling of ROSMAP brain tissues were submitted to the AD Knowledge Portal in two rounds. Round 1 (submitted in 2018) provides data from 400 individuals. Round 2 (submitted in 2022) provides data from an additional 210 individuals. | https://www.synapse.org/#!Synapse:syn21266454 |
| ROSMAP_RNAseq_FPKM_gene | Samples were extracted using Qiagen's miRNeasy mini kit (cat. no. 217004) and the RNase free DNase Set (cat. no. 79254), and quantified by Nanodrop and quality was evaluated by Agilent Bioanalyzer. | https://www.synapse.org/#!Synapse:syn3505720 |
| ROSMAP.CNV.Matrix(Mutation) | The TCGA Unified Ensemble "MC3" gene-level mutation dataset identifies somatic mutations in various cancers, marking non-silent mutations (1) that alter protein sequences and wild type (0) for no mutations. | https://www.synapse.org/#!Synapse:syn26263118 |
| GEO GPL16304 Platform | Illumina HumanMethylation450 BeadChip [UBC enhanced annotation v1.0] | https://www.ncbi.nlm.nih.gov/geo/query/acc.cgi?acc=GPL16304 |
| ROSMAP_clinical | Contains patient clinical features. A large amount of clinical and pathological data have been collected from individuals in the ROSMAP studies. The remainder of the clinical and pathological data may be accessed directly from the Rush Alzheimer's Disease Center. | https://www.synapse.org/#!Synapse:syn3191087 |

#### KEGG regulatory network

Genes for constructing the knowledge graph were selected by intersecting multi-omics datasets with gene regulatory networks from the KEGG database, which comprises 2121 genes, 19751 protein-protein interactions and 26114 edges for UCSC Xena dataset; 2146 genes, 19867 protein-protein interactions and 26305 edges. After this intersection, the resulting number of entities in constructed biomedical knowledge graph was 8484 and 8584 for UCSC Xena and ROSMAP respectively.

### 2.2. The mosGraphGPT model

#### Problem Formulation

The overall architecture of mosGraphGPT model was demonstrated in **Figure 1**. Given the bulk-seq multi-omics datasets of 𝒳^(Epi)^ for epigenomics, 𝒳^(Geno)^ for genomics, 𝒳^(Tran)^ for transcriptomics and 𝒳^(Prot)^ for proteomics and clinical dataset 𝒴^(*c*)^, the integration over biomedical knowledge graph was completed by mosGraphGen^2^ with 𝒢= (*V, E*), which can be decomposed into subgraphs 𝒢_int_ = (*V*_int_, *E*_int_), where |*V*_int_| = *n*_Epi_ + *n*_Geno_ + *n*_Tran_ + *n*_Prot_ = *n* = |*V*| and 𝒢_PPI_ = (*V*_PPI_, *E*_PPI_), where |*V*_PPI_| = *n*_Prot_ and 𝒢_int_ = 𝒢\ 𝒢_PPI_. Correspondingly, adjacency matrix *A* ∈ ℝ^*n*×*n*^ for whole graph 𝒢 will be generated and adjacency matrix *A*_int_ ∈ ℝ^*n*×*n*^ for internal signaling flows from promoters to proteins via central dogma. What’s more, protein-protein interactions will be represented by adjacency matrix *A*_PPI_ ∈ ℝ^*n*×*n*^ and *A* = *A*_int_ + *A*_PPI_. Furthermore, patient multi-omics feature 𝒳 = {*X*^(1)^, *X*^(2)^, …, *X*^(*m*)^, …, *X*^(*M*)^} will be generated, where *X*^(*m*)^ ∈ ℝ^*n*×*d*^, *d* equals the number of multi-omics data features and *n* equals the number of nodes. With above processed pretraining datasets, the encoder model, *f*_pre_(⋅), will be pretrained by self-supervised learning by reconstructing edges and degree of nodes. Similarly, the input bulk-seq multi-omics datasets with clinical features can also be generated with 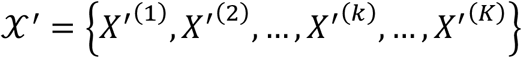 and 𝒴′ = {*y*′^(1)^, *y*^′(2)^, …, *y*^′(*k*)^, …, *y*′^(*K*)^} (*X*′^(*k*)^ ∈ ℝ^*n*×*d*^, 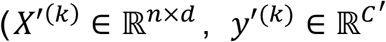 and *C*^′^ is the number of patient types). Regarding the biomedical knowledge graph, internal signaling flows graph, protein-protein interactions graph, all of them share the same network structures with 𝒢, 𝒢_int_, 𝒢_PPI_ and adjacency matrices *A, A*_int_, *A*_PPI_. Hence, the similar graph structures were generated by 𝒢′, 𝒢′_int_, 𝒢′_PPI_ and adjacency matrices *A*′, *A*′_int_, *A*′_PPI_ .And our proposed model can be denoted as *f*(⋅) to predict the patient types with 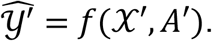.

**Figure 1.**
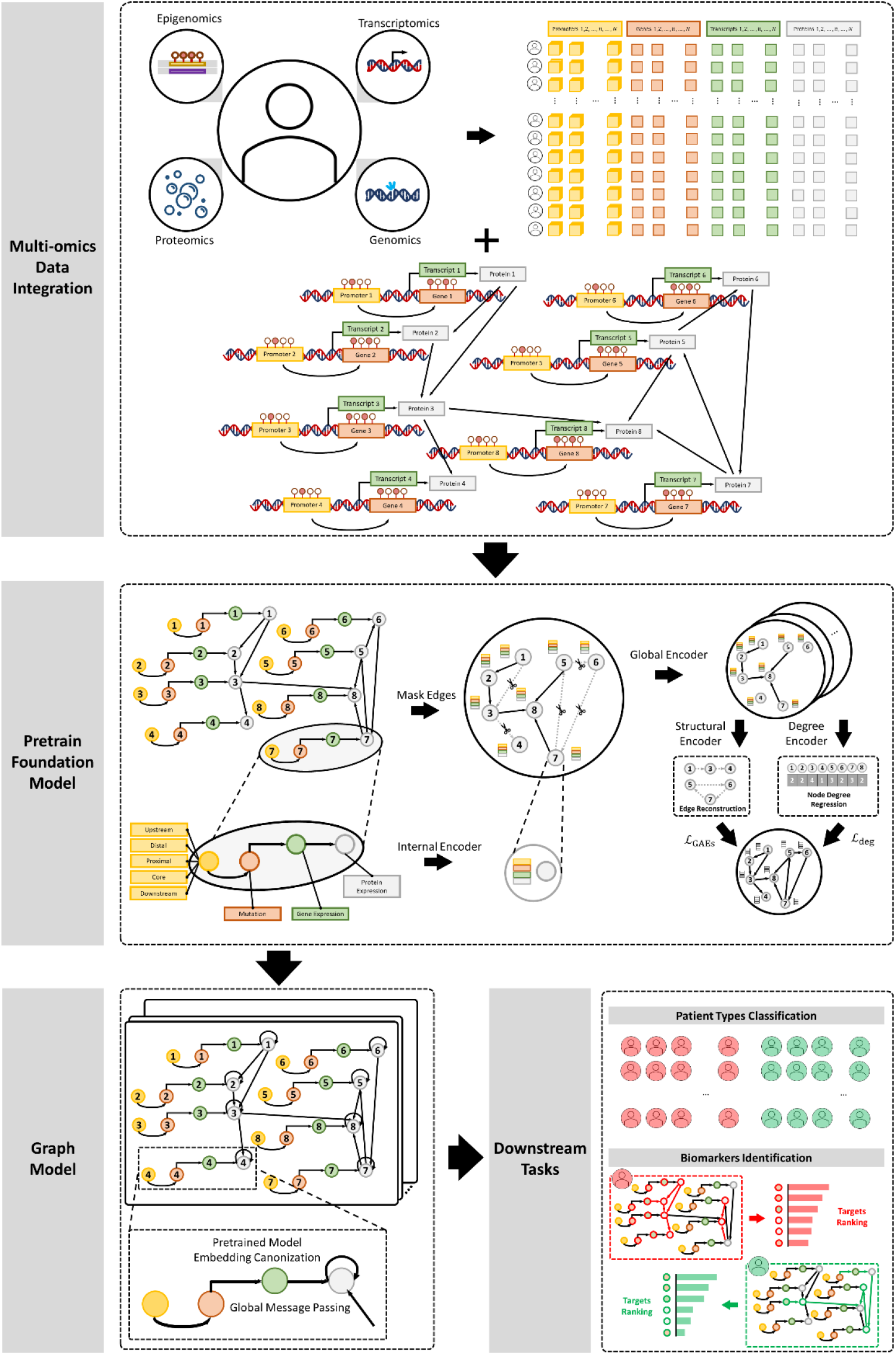
Architecture of ***mosGraphGPT***.

#### Pretrain Foundation Model

During the pretraining stage, the constructed biomedical knowledge graph *A* can be decomposed into 2 subgraph paths *A*_int_ and *A*_PPI_, where message propagation will also be separated with 2 steps for internal signaling flows from promoters to proteins and protein signaling flows between protein-protein interactions. Since, it is the protein-protein interactions that the mosGraphGPT model would like to reconstruct, earlier stage message passing was accomplished to propagate information to the protein nodes with:

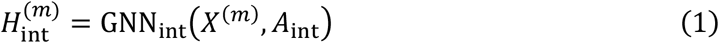

where 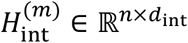 is the node features which was diffused from epigenomics, genomics, transcriptomics and proteomics onto the ending protein nodes and GNN_int_ is the internal signaling flows encoder. Meanwhile, to mask the edges for pretrain model to reconstruct, the random masking function Γ will be generated following a specific distribution, e.g., Bernoulli distribution:

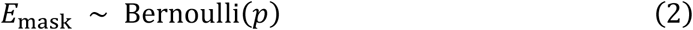

where *p* < 1 is the ratio of the masked edges for the protein-protein interactions graph 𝒢_PPI_. Hence, the masked protein-protein interactions graph will be denoted as 𝒢_mask_ and the unmasked or visual protein-protein interactions graph will be denoted as 𝒢_vis_, where 𝒢_PPI_ = 𝒢_vis_ ⋃ 𝒢_mask_. Correspondingly, the masked adjacency matrix *A*_mask_ and visual adjacency matrix *A*_vis_ will be generated by the masking function Γ. Afterwards, the global encoder will be used to generate the node embeddings by

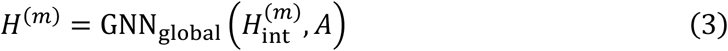

where 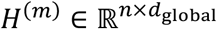 and GNN_global_ is the graph neural network message propagation for signaling flows in global paths. With the global node embeddings, structural decoder and degree decoder were built to learn the pretrain model. In details, the structural decoder *u*_ω_ with parameters ω will decode to probability of the edge connection between node *p* and *q* by

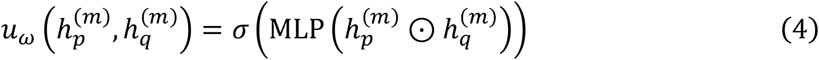

where 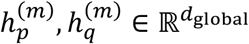 are global node embedding for node *p* and *q* for patient *m*; MLP is the multilayer perceptron and ⊙ is the element-wise product. Moreover, the degree decoder *ν*_*ϕ*_ was constructed by

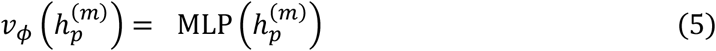

where *ϕ* is the parameter learnt from the degree decoder and the decoder aims to reconstruct the node degree with regression.

In sum, the pretrain model reconstruction loss, ℒ^(*m*)^, will be calculated by edge reconstruction loss, which measures how well the model can rebuild the edge connect in the protein-protein interactions network and degree regression loss, which measures how closely the prediction of the node degree matches the degree of nodes in original graph 𝒢 with

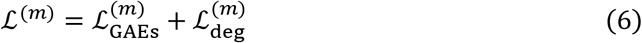

where edge reconstruction loss, ℒ_GAEs_, is self-supervised learning objective loss by optimizing the cross-entropy loss via

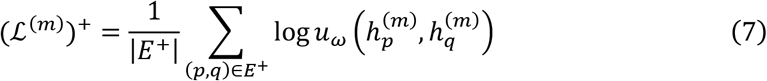

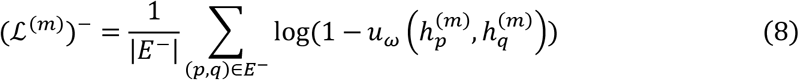

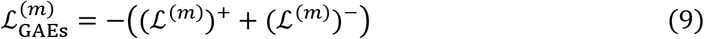

where *E*^+^ is a set of positive edges while *E*^−^ is a set of negative edges sampled from the protein-protein interaction graph 𝒢_PPI_ and degree reconstruction loss will be calculated with mean squared error (MSE) between the original degree of nodes and the predicted ones

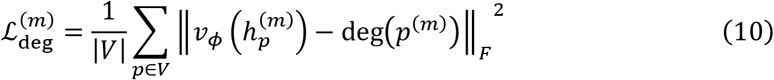

where deg(⋅) is the degree function which can generate the degree of node *p* for patient *m* over whole graph 𝒢.

#### Graph Model Construction

With the pretrained encoder function 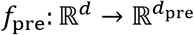 composed of internal encoder and global encoder shown in formula (1) and (3), the input feature can generate embedding 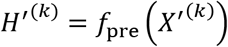 which can be used as the graph canonization via residual process with

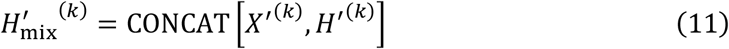

where 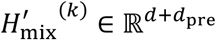 is the concatenated node features for patient *k*. To predict the patient *k* outcome, the global message propagation was conducted via

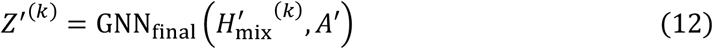

where 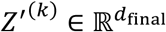 and transformer-based message passing network was leveraged here as the GNN_final_. The cross-entropy (CE) function was used via

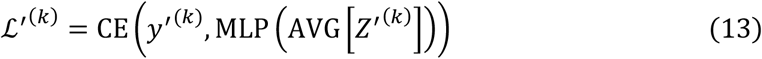

where mean aggregation pooling function AVG in PyTorch and linear transformation with MLP was leverage.

### 2.3. Downstream Tasks

#### Predict patient types

Given the embedded features 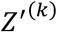 for patient *k*, the prediction of the patient type can be generated by

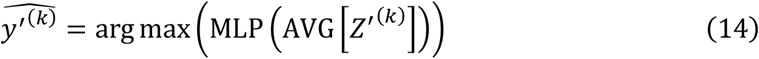

where 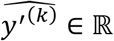.

#### Biomarkers Identification via Attention

Based on the attention extracted from transformer, the weighted adjacency matrix 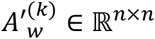 will be generated. Furthermore, the average weighted adjacency matrix of specific patient type *c* can be calculated by

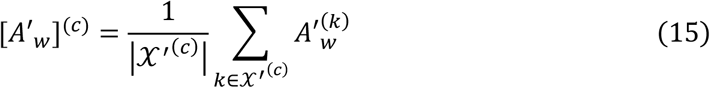

where 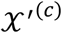 is the set of specific patient type *c*.

## 3. Results

### 3.1. mosGraphGPT model results

#### Experimental settings

The UCSC xena multi-omics dataset was used as the pretrain model, which contains 3592 cancer patients with 2121 genes, 8484 node entities, 19751 protein-protein interactions and 26114 relations. Early stopping strategies was employed for self-supervised pretraining process. Afterwards, for training the whole model, ROSMAP was loaded with 128 samples with 2146 genes, 8584 node entities, 19867 protein-protein interactions and 26305 relations. Specifically, the 5 fold cross validation was used to train and test the proposed model, *mosGraphGPT*. To evaluate the model performance in terms of synergy score prediction for drug combinations, we conducted 5-fold cross validation. The *mosGraphGPT* model was implemented using PyTorch and PyTorch Geometric, with the Adam optimizer of setting weight decay as 1 × 10^−20^ and ϵ as 1 × 10^−7^ employed for training.

#### Model performance and comparison

The average prediction accuracy was about **75.09%** the test data on ROSMAP AD dataset. The results indicated the feasibility of patient outcome prediction using a graph neural network with a small set of core signaling pathways genes. Moreover, the proposed model was compared with other graph neural networks (see **Table 3**), which included the GNN model with mean aggregation in transductive mode^38^ (no sampling for neighborhood function) and Graph Attention network^39^ (GAT), Graph Isomorphism Network (GIN)^40^ and UniMP^41^.

**Table 3.**
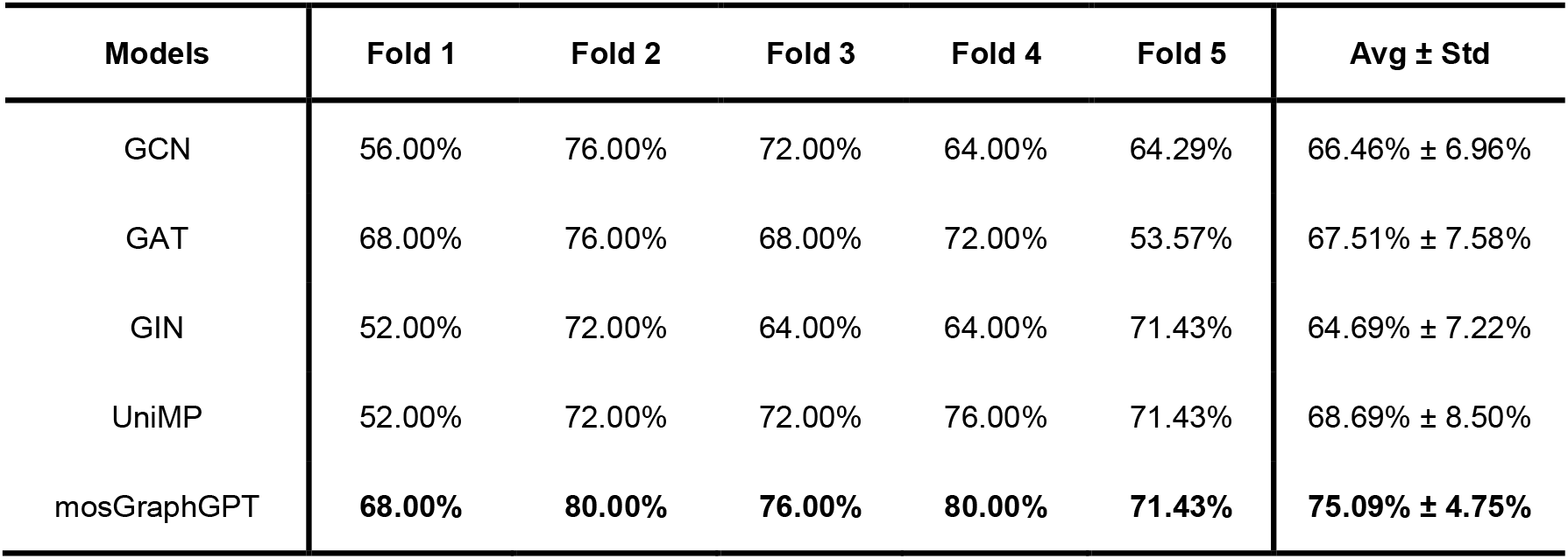
Model comparison with other GNN networks (AD vs. Non-AD)

| Models | Fold 1 | Fold 2 | Fold 3 | Fold 4 | Fold 5 | Avg ± Std |
| --- | --- | --- | --- | --- | --- | --- |
| GCN | 56.00% | 76.00% | 72.00% | 64.00% | 64.29% | 66.46% ± 6.96% |
| GAT | 68.00% | 76.00% | 68.00% | 72.00% | 53.57% | 67.51% ± 7.58% |
| GIN | 52.00% | 72.00% | 64.00% | 64.00% | 71.43% | 64.69% ± 7.22% |
| UniMP | 52.00% | 72.00% | 72.00% | 76.00% | 71.43% | 68.69% ± 8.50% |
| mosGraphGPT | **68.00%** | **80.00%** | **76.00%** | **80.00%** | **71.43%** | **75.09% ± 4.75%** |

### 3.2. Downstream tasks

To investigate the potential MoS, a core signaling subnetwork was generated based on the trained model. Specifically, the integrated signaling flow networks were obtained from the large signaling network based on the averaged trained directional weight matrices on the 5 splits of test datasets on ROSMAP-AD. With averaging the attention weight adjacency matrices 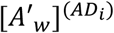 and 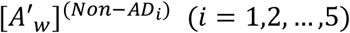 on each fold, the weight adjacency matrices [*A*′_*w*_]^(*AD*)^ for AD samples and [*A*′_*w*_]^(*Non*−*AD*)^ for Non-AD samples are generated. Afterwards, the cell line specific gene degrees will be calculated based on following formula:

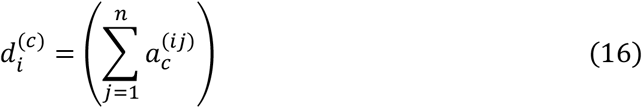

where 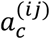 is the element in the *i*-th row and *j*-th column of the matrices [*A*′_*w*_]^(*AD*)^ for AD samples (*c* = *AD*) and [*A*′_*w*_]^(*Non*−*AD*)^ for Non-AD patients (*c* = *Non* − *AD*), which measures the link strength between node *i* and node *j*. Hence, 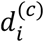 is the weighted degree for node *i* from specific sample type *c*.

Afterwards, the unimportant signaling flows in the attention-based matrix for certain type of patient will be filtered out by

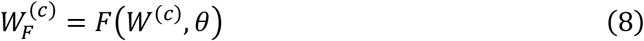

where *F*(⋅) is the filtering mapping function by providing selection of each element in the matrix with

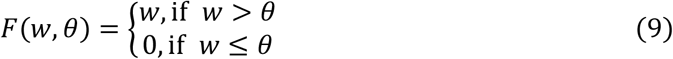

where *w* ∈ ℝ is the element in the input matrix and 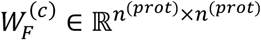 is the filtered matrix. Hence, the filtered node set for patient type *c*, 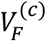, will be generated by removing independent nodes and nodes in those small connected components with number of nodes lower than *ϕ*, resulting in 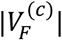 nodes.

Subsequently, p-values for the gene features, such as methylations in promoter nodes, mutations and genes expression in gene nodes and proteins expression in protein nodes were calculated. The p-value calculation for these features was conducted by using the chi-squared test to check the differences between AD/non-AD samples or female/male of AD patients. This statistical method determined whether there were significant differences in the gene features between the samples of AD/non-AD or female/male from AD. By constructing contingency tables and performing the chi-squared test for each gene feature, p-values indicating the statistical significance of the observed differences were obtained. Ultimately, the top *T* gene features associated with AD or gender were selected based on these p-values.

After finalized important gene features ranked by p-values in top *T*, the network was pruned by iteratively removing the nodes which are only connected to one another unimportant node in a linear branch with node recursive algorithm (check details of this algorithm in **Appendix A** and **Figure S1**). This ensures that each remaining nodes is either linked to an important node or is part of a more complex interaction network, enhancing the purity and reliability of the gene interaction data.

Subsequently, nodes degree were calculated to identify hub node (node degree larger than 2). The set of middle nodes for certain path *t* which connects two hub nodes *u* and *ν* can be denoted as 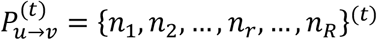, where *λ* + 1 is the length of path. Hence, the average edge weight on the path 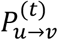 can be generated by

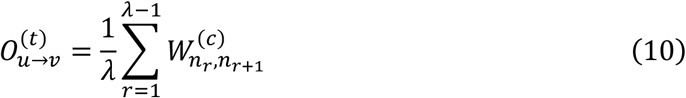

where 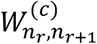 is the edge weight from node *n*_*r*_ to node *n*_*r*+1_. For all of the paths detected between the hub node *u* and hub node *ν*, the nodes on the top *β* paths will be kept. Additionally, p-value middle nodes, which are crucial due to their statistical significance, will be retained along with middle nodes that are adjacent to these p-value nodes. (check **Appendix Section A.2** for details).

#### Uncovering AD associated targets and signaling pathways

Setting the edge threshold *θ* as 0.15 and filtering out small components with nodes fewer than 25 (*ϕ*=25), the core signaling network flows for AD and non-AD samples were generated. The AD core signaling network included 335 potential important protein nodes, and the non-AD network contained 408 potential important protein nodes. To further discover the top 100 important node entities in the whole network, p-value was leveraged to measure the importance of the nodes in this core signaling network. Visualization of the core signaling networks were also accomplished with the pruning algorithm (check **Appendix A**) to mark the important nodes in the network (shown in **Figures 2-4**). Through the polished algorithm, the AD network was refined to 256 protein nodes and the non-AD network to 267 protein nodes. This significantly reduced the impact of irrelevant nodes on the core signaling networks visualization, providing a clearer depiction of protein-protein interactions.

**Figure 2.**
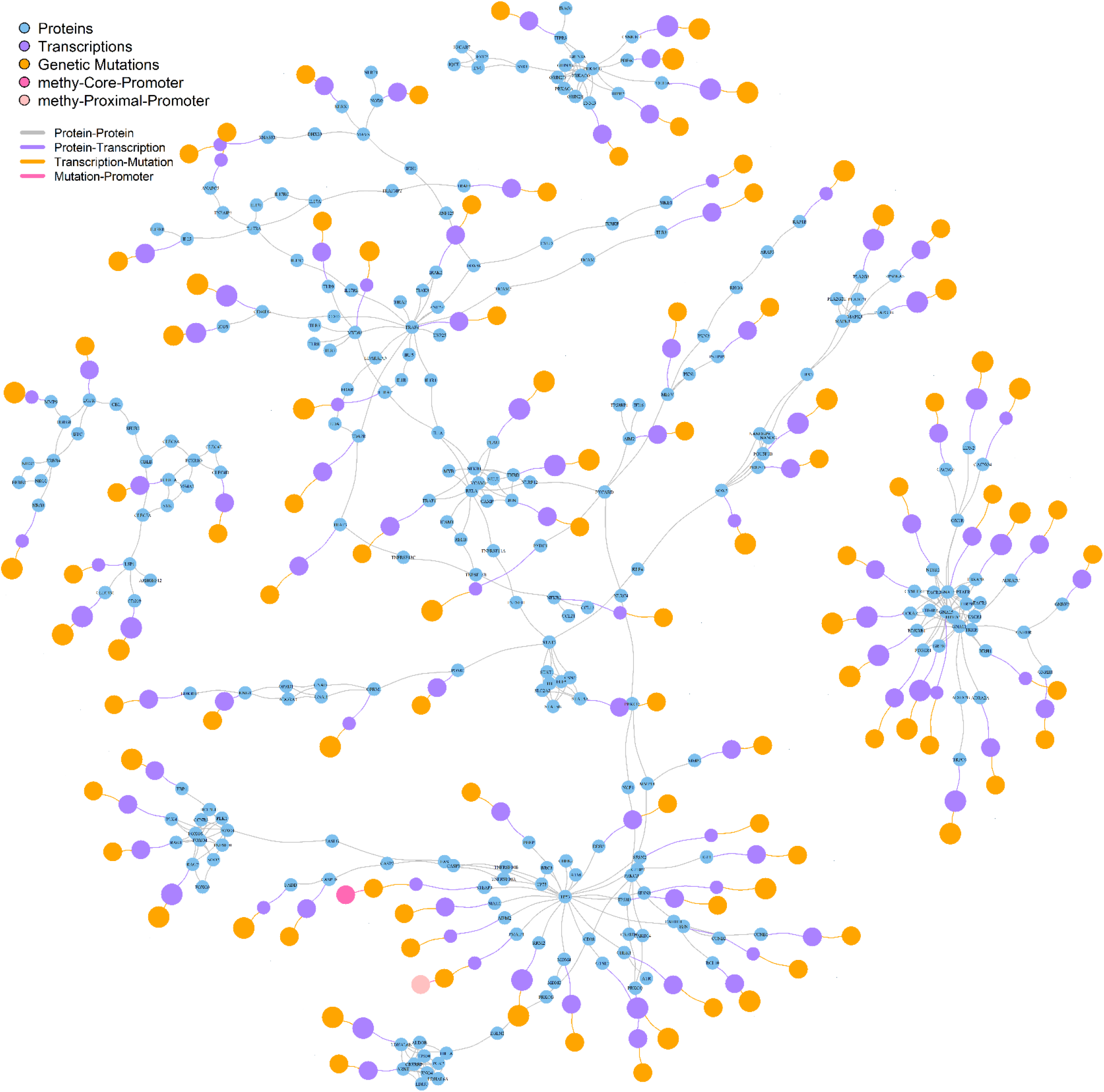
Important signaling network flows for AD patients with top 100 gene features ranked by p-values

**Figure 3.**
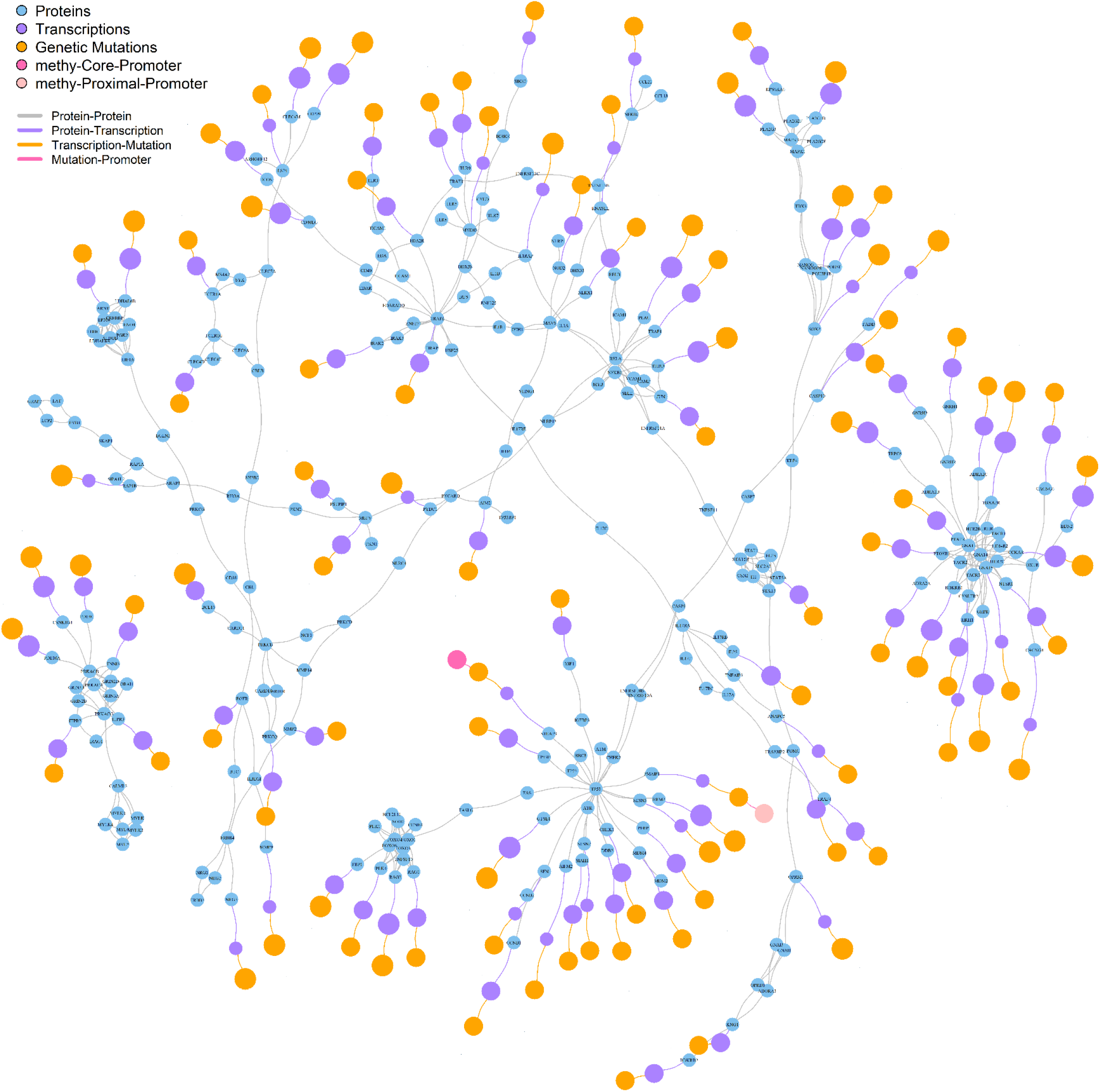
Important signaling network flows for non-AD patients with top 100 gene features ranked by p-values

**Figure 4.**
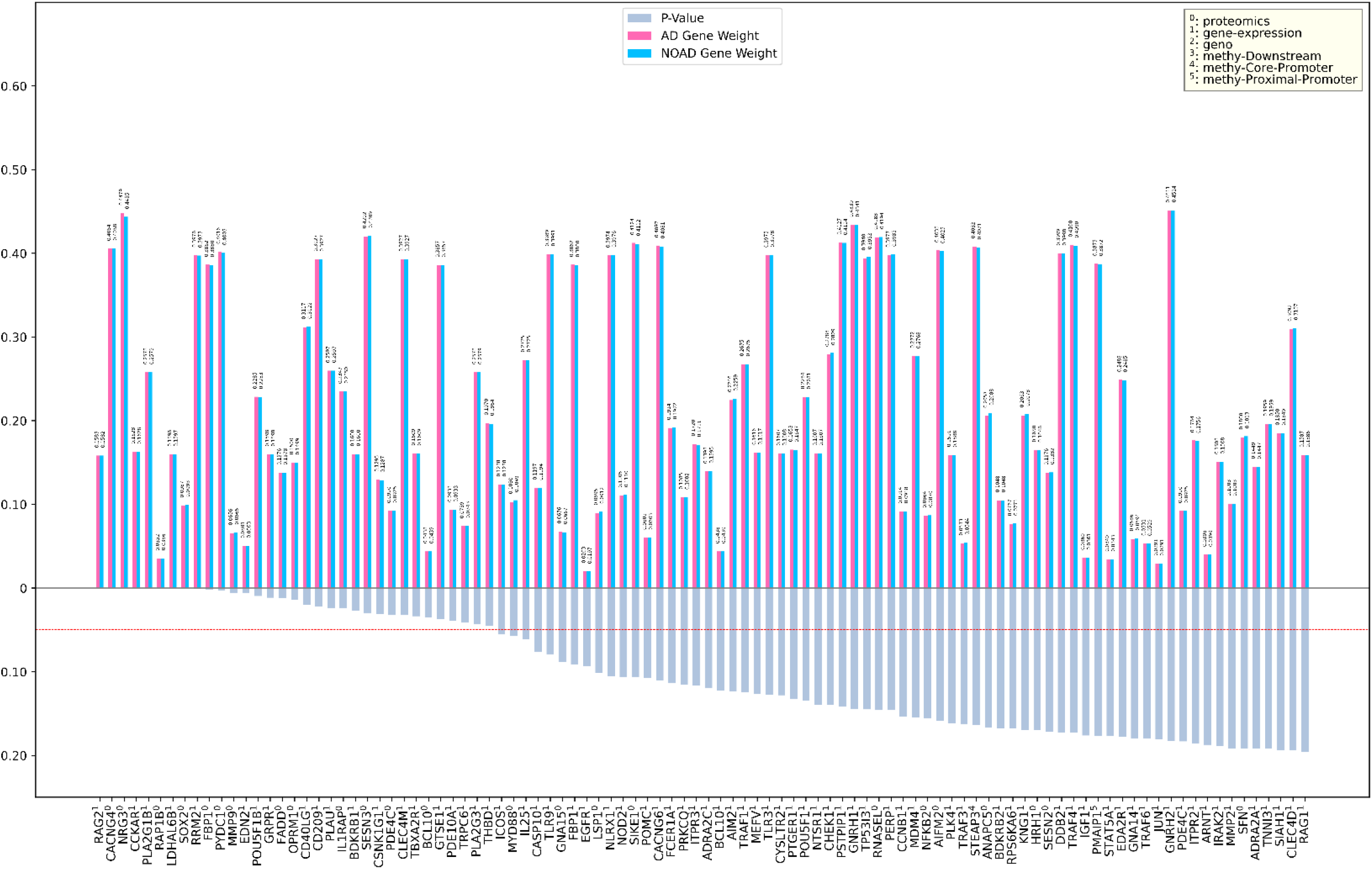
Top 100 important gene features for AD samples ranked by p-value

### 3.3. Validation on identified biomarkers

Based on the top 100 gene features, which include promoters, mutations, transcriptions, and proteins, selected for Alzheimer’s disease (AD) samples through the use of p-values, a comprehensive validation process has been undertaken. This selection was made to identify the most statistically significant gene features relevant to AD, thereby enabling a focused analysis of the underlying biological mechanisms. The subsequent validation involved conducting a pathway enrichment analysis, which is a critical step in understanding the functional implications of these gene features. This analysis helps to identify biological pathways that are significantly enriched in the dataset, providing insights into the molecular processes and pathways that may be disrupted in AD. The results of this pathway enrichment analysis are presented in **Figures 5-6** and detailed in **Table 4**. These findings are essential for corroborating the initial selection of gene features and for highlighting potential targets for further investigation. The integration of these results into the broader context of AD research underscores the importance of pathway enrichment analysis in validating genetic and molecular data, thus contributing to a more nuanced understanding of the disease’s pathogenesis and potential therapeutic avenues.

**Figure 5.**
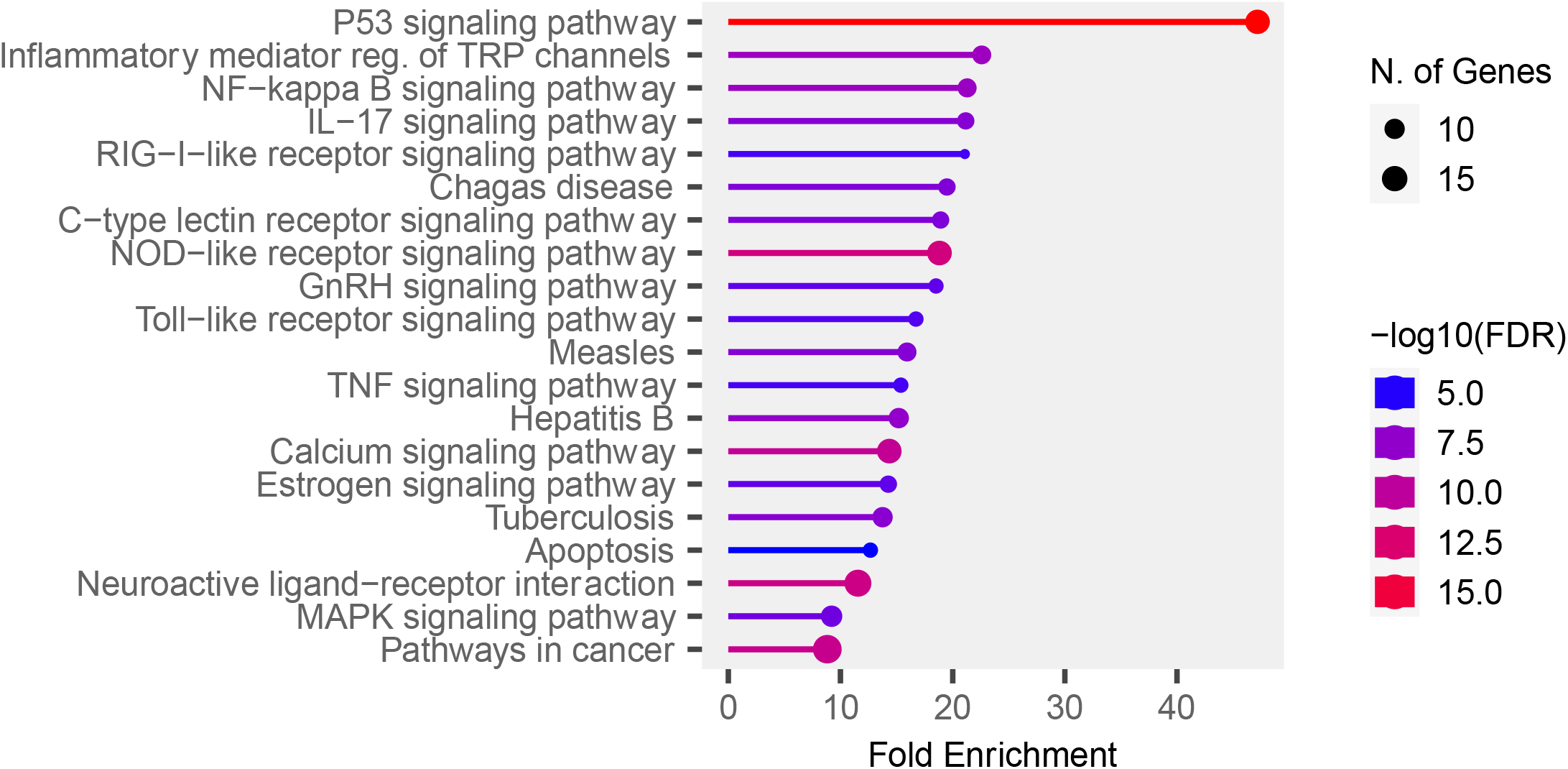
Top 20 pathways lollipop plot

**Figure 6.**
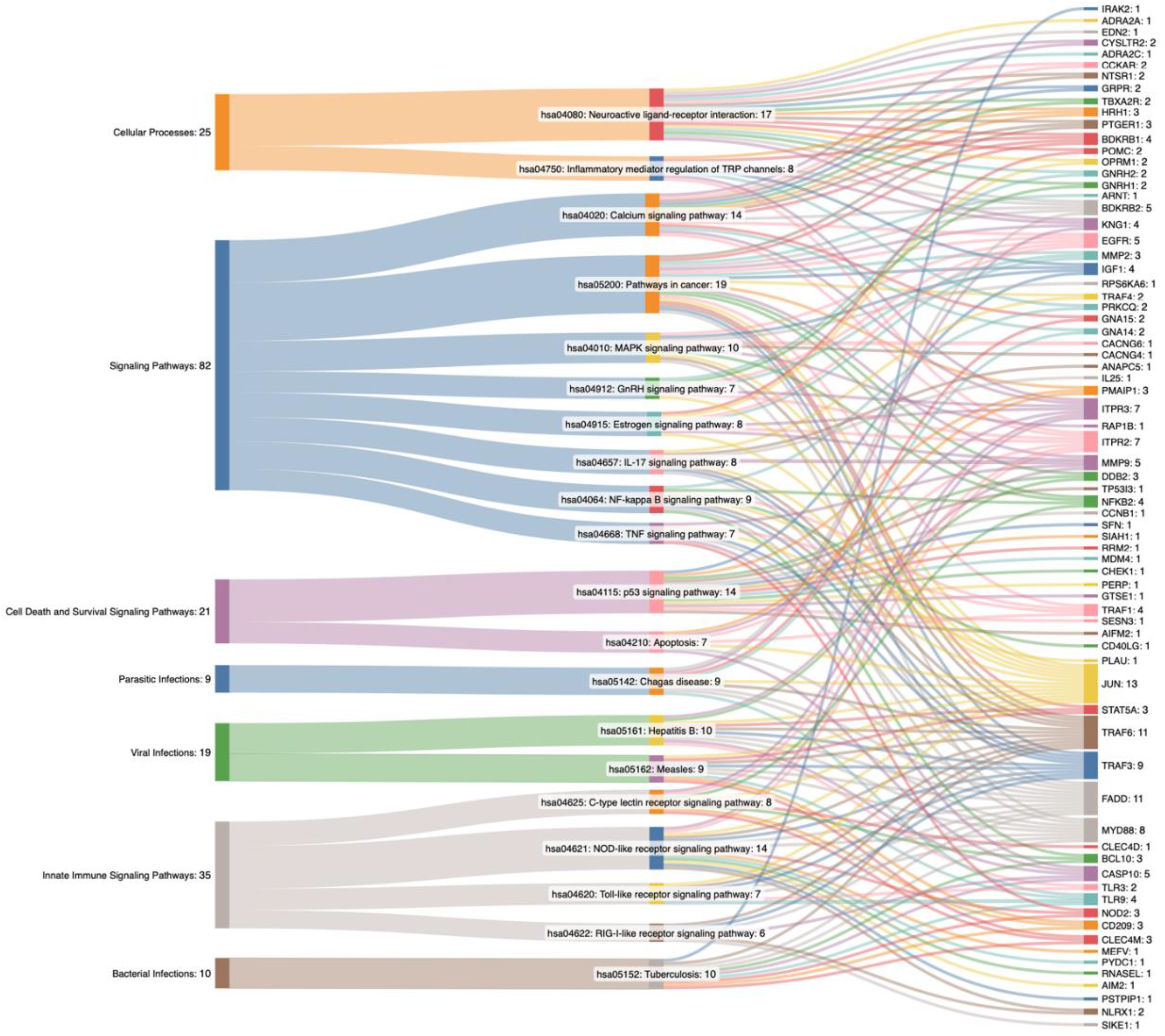
Top 20 pathways Sankey chart

**Table 4.** Pathway enrichment analysis for top 20 gene features ranked by p-values.

| Type | Pathway | Number of Genes | Genes | P-Value | FDR |
| --- | --- | --- | --- | --- | --- |
| Signaling Pathways | hsa05200: Pathways in cancer | 19 | BDKRB1,MMP2,DDB2,TRAF4,KNG1,EGFR,PTGER1,FADD,IGF1 | 6.1E-12 | 4.1E-10 |
|  |  |  | ,PMAIP1,STAT5A,ARNT,NFKB2,<br>JUN,MMP9,TRAF1,TRAF6,BDK<br>RB2,TRAF3 |  |  |
| Cellular<br>Processes | hsa04080: Neuroactive<br>ligand-receptor<br>interaction | 17 | BDKRB1,GNRH2,KNG1,GNRH1,<br>ADRA2A,CYSLTR2,PTGER1,CC<br>KAR,NTSR1,EDN2,GRPR,HRH1<br>,POMC,ADRA2C,TBXA2R,OPR<br>M1,BDKRB2 | 5.42E-13 | 6.07E-11 |
| Cell Death and<br>Survival<br>Signalig<br>Pathways | hsa04115: p53 signaling<br>pathway | 14 | TP53I3,CCNB1,DDB2,IGF1,PMA<br>IP1,SFN,SIAH1,RRM2,MDM4,C<br>HEK1,PERP,GTSE1,SESN3,AIF<br>M2 | 3.49E-18 | 1.17E-15 |
| Innate Immune<br>Signalig<br>Pathways | hsa04621: NOD-like<br>receptor signaling<br>pathway | 14 | MEFV,NOD2,FADD,PYDC1,RNA<br>SEL,AIM2,JUN,ITPR3,ITPR2,NL<br>RX1,MYD88,TRAF6,PSTPIP1,T<br>RAF3 | 2.52E-13 | 4.23E-11 |
| Signaling<br>Pathways | hsa04020: Calcium<br>signaling pathway | 14 | BDKRB1,GNA15,EGFR,CYSLTR<br>2,PTGER1,CCKAR,NTSR1,ITPR<br>3,GNA14,GRPR,ITPR2,HRH1,TB<br>XA2R,BDKRB2 | 9.45E-13 | 7.94E-11 |
| Viral Infections | hsa05161: Hepatitis B | 10 | DDB2,CASP10,TLR3,FADD,STA<br>T5A,JUN,MMP9,MYD88,TRAF6,<br>TRAF3 | 7.38E-09 | 3.1E-07 |
| Bacterial<br>Infections | hsa05152: Tuberculosis | 10 | IRAK2,CASP10,NOD2,FADD,CD<br>209,CLEC4M,TLR9,BCL10,MYD<br>88,TRAF6 | 1.21E-08 | 4.53E-07 |
| Signaling Pathways | hsa04010: MAPK signaling pathway | 10 | RAP1B,CACNG6,CACNG4,RPS6KA6,EGFR,IGF1,NFKB2,JUN,MYD88,TRAF6 | 1.47E-06 | 2.9E-05 |
| Parasitic Infections | hsa05142: Chagas disease | 9 | GNA15,KNG1,FADD,TLR9,JUN,GNA14,MYD88,TRAF6,BDKRB2 | 2.13E-09 | 1.19E-07 |
| Signaling Pathways | hsa04064: NF-kappa B signaling pathway | 9 | PRKCQ,NFKB2,BCL10,CD40LG,PLAU,TRAF1,MYD88,TRAF6,TRAF3 | 2.51E-09 | 1.2E-07 |
| Viral Infections | hsa05162: Measles | 9 | FADD,CD209,CLEC4M,STAT5A,TLR9,JUN,MYD88,TRAF6,TRAF3 | 3.22E-08 | 9.02E-07 |
| Signaling Pathways | hsa04657: IL-17 signaling pathway | 8 | ANAPC5,TRAF4,FADD,IL25,JUN,MMP9,TRAF6,TRAF3 | 2.41E-08 | 8.09E-07 |
| Cellular Processes | hsa04750: Inflammatory mediator regulation of TRP channels | 8 | BDKRB1,PRKCQ,KNG1,IGF1,ITPR3,ITPR2,HRH1,BDKRB2 | 2.81E-08 | 8.6E-07 |
| Innate Immune Signaling Pathways | hsa04625: C-type lectin receptor signaling pathway | 8 | CLEC4D,CD209,CLEC4M,NFKB2,BCL10,JUN,ITPR3,ITPR2 | 5.1E-08 | 1.32E-06 |
| Signaling Pathways | hsa04915: Estrogen signaling pathway | 8 | MMP2,EGFR,JUN,MMP9,ITPR3,ITPR2,POMC,OPRM1 | 3.5E-07 | 8.39E-06 |
| Signaling Pathways | hsa04912: GnRH signaling pathway | 7 | MMP2,GNRH2,EGFR,GNRH1,JUN,ITPR3,ITPR2 | 3.57E-07 | 8.39E-06 |
| Innate Immune Signaling Pathways | hsa04620: Toll-like receptor signaling pathway | 7 | TLR3,FADD,TLR9,JUN,MYD88,TRAF6,TRAF3 | 7.97E-07 | 1.67E-05 |
| Signaling<br>Pathways | hsa04668: TNF<br>signaling pathway | 7 | CASP10,NOD2,FADD,JUN,MMP<br>9,TRAF1,TRAF3 | 1.54E-06 | 2.9E-05 |
| Cell Death and<br>Survival<br>Signaling<br>Pathways | hsa04210: Apoptosis | 7 | CASP10,FADD,PMAIP1,JUN,TR<br>AF1,ITPR3,ITPR2 | 4.33E-06 | 7.28E-05 |

#### Cell Death and Survival Signaling

As a crucial transcription factor, p53 regulates DNA repair, cell cycle control, apoptosis, and oxidative stress response. In AD, p53 function is notably disrupted, leading to increased DNA damage and impaired repair mechanisms. Elevated phosphorylated p53 levels and altered oligomerization states in AD patients’ temporal lobes indicate compromised DNA damage response and repair capabilities^42^. p53 also forms oligomers and fibrils that interact with tau oligomers, potentially seeding further p53 aggregation and mislocalization outside the nucleus, impairing its nuclear functions^43^. This aggregation and tau interaction may contribute to lethal cell cycle re-entry and abnormal cell death in AD. Additionally, p53 signaling intersects with other dysregulated pathways in AD, such as WNT and NFkB, particularly in inhibitory neurons, where decreased p53 activity and altered transcription factor activity are observed^44^. The complexity of p53’s role in AD is heightened by its regulation through post-translational modifications, affecting its conformation and function, and potentially influencing amyloid and tau pathways^45^. Oxidative stress in AD patients exacerbates p53 dysfunction, as indicated by increased protein carbonylation and impaired cGAS-STING-interferon signaling, crucial for immune-stimulated DNA repair^42^.

Apoptosis is a programmed cell death mechanism essential for maintaining cellular homeostasis and regulating cell turnover, but its dysregulation can lead to neurodegenerative disorders, including AD^46,47^. In AD, several pathological features such as Aβ plaques, hyperphosphorylated tau tangles, inflammation, mitochondrial dysfunction, and oxidative stress trigger an abnormal apoptotic cascade in critical brain regions like the cerebral cortex and hippocampus^46^. This cascade involves various molecular pathways, including PI3K/AKT, JNK, MAPK, and mTOR signaling, which ultimately result in neuronal death and correlate with the severity of dementia^46^. Additionally, apoptosis interacts with necroptosis, another form of programmed cell death, which is also activated in AD brains and contributes to neuroinflammation and neuronal death^48–50^. The interplay between apoptosis and necroptosis exacerbates the neurodegenerative process, as necroptosis can be triggered by factors such as hyperglycemia and reactive oxygen species, which are prevalent in AD. Therapeutic strategies targeting apoptotic pathways, such as caspases and other apoptotic regulators, have been explored to mitigate neuronal loss and slow disease progression^46,47^.

#### Signaling Pathways

NF-κβ signaling is central to neuroinflammation and oxidative stress, exacerbating neurodegeneration by interacting with reactive microglia, astrocytes, and various molecular factors, while also influencing amyloid plaque clearance and neuronal survival^51^. Dysregulated calcium signaling disrupts neuronal function and survival by causing mitochondrial failure, oxidative stress, and chronic neuroinflammation, leading to NFTs and Aβ plaques^52^. The ER-mitochondria membrane contact site is particularly critical for calcium homeostasis, and its disruption further exacerbates AD pathology^53^. Estrogen signaling, particularly in women, plays a multifaceted role in AD, with estrogen deficiency post-menopause promoting amyloid precursor protein processing into senile plaques and increasing tau phosphorylation^54^. Estrogen also affects glucose metabolism and WNT signaling, contributing to neuropathology. The interaction between estrogen and APOE genotype modulates AD risk, with estrogen receptors’ reduced activity accelerating disease progression^55^.

GnRH, along with luteinizing hormone (LH) and activins, possesses neuronal receptors that are distributed throughout the limbic system, which is crucial for cognitive functions and is notably affected in AD^56^. Dysregulation of the HPG axis during menopause and andropause leads to elevated levels of GnRH and LH, while sex steroid signaling decreases, potentially promoting neurodegenerative changes^56^. Elevated LH levels, which are a consequence of this dysregulation, have been implicated in the amyloidogenic processing of APP^56,57^. Furthermore, LH is known to cross the blood-brain barrier and its receptors are highly concentrated in the hippocampus, a region particularly vulnerable to AD^56^. Pharmacological interventions that suppress LH release, such as leuprolide acetate, have shown promise in reducing Aβ deposition and improving cognitive performance in animal models of AD, suggesting a potential therapeutic avenue^56,57^. Epidemiological data also support this connection, as reduced neurodegenerative disease incidence has been observed among prostate cancer patients treated with GnRH agonists, which lower LH levels^56^.

Dysregulation of MAPK signaling, particularly through the ERK/MAPK1 pathway, has been implicated in the development of AD pathogenesis^58^. Specifically, phosphorylated ERK (p-ERK) has been identified as a critical regulator of pro-inflammatory activation of microglia, which are immune cells in the brain that contribute to neuroinflammation in AD^59,60^. This pro-inflammatory state is further exacerbated by the JAK/STAT signaling pathway, which is activated by overactive microglia and astrocytes, leading to a chronic neuroinflammatory environment that is characteristic of AD. Additionally, the PI3K-Akt pathway, which interacts with MAPK signaling, is involved in regulating cell survival and metabolic functions, and its dysregulation is linked to Aβ and NFTs^61^. The complex interplay between these pathways is evident as the PKR/P38/RIPK1 signaling axis, part of the stress-activated MAPK pathway, is highly activated in AD brains, leading to Aβ accumulation, tau phosphorylation, and cognitive decline^62^. Experimental models have shown that modulating miRNAs that regulate MAPK signaling can improve cognitive deficits, highlighting the therapeutic potential of targeting this pathway^61^.

In AD, TNF signaling has been implicated in promoting necroptosis. This is evidenced by increased expression of necroptosis-related proteins such as phosphorylated RIPK3 and MLKL in the AD brain, particularly in CA1 pyramidal neurons, which correlates inversely with neuron density^63^. Additionally, TNF exposure in human iPSC-derived neurons increases necroptotic cell death, which can be mitigated by inhibitors targeting RIPK1, RIPK3, and MLKL, suggesting potential therapeutic intervention points^63^. Furthermore, TNF-mediated neuroinflammation is exacerbated by the interaction of misfolded proteins with pattern recognition receptors on astroglia and microglia, leading to the release of inflammatory mediators that contribute to disease progression^64^. Genome-wide analyses have identified several genes associated with sporadic AD that control inflammatory responses and glial clearance of misfolded proteins, highlighting the critical role of immune processes in AD pathogenesis^64^. Interestingly, patients with rheumatoid arthritis and other systemic inflammatory diseases treated with TNF-α blocking agents show a reduced probability of developing dementia, suggesting that TNF-α inhibition could be a viable strategy for preventing AD and preserving cognitive function^65^.

Studies have shown that IL-17A levels are elevated in the brains of AD patients and animal models, suggesting its involvement in disease progression^66,67^. When IL-7A is overexpressed, worsening of cognitive functions is observed^67^. IL-17A also exacerbates neuroinflammation by facilitating the infiltration of immune cells such as CD8+ T lymphocytes and myeloid cells into the brain, which in turn accelerates the production of pro-inflammatory chemokines like CXCL1 and CXCL9/10 by glial cells^67^. This inflammatory milieu promotes Aβ accumulation and synaptic dysfunction, leading to cognitive deficits. Furthermore, IL-17A has been shown to induce neural damage directly when administered to primary hippocampal neurons, and its inhibition via neutralizing antibodies can ameliorate Aβ-induced neurotoxicity and cognitive decline by downregulating the TRAF6/NF-κB pathway^66^. Additionally, the depletion of gut bacteria, which reduces IL-17A-expressing T cells, has been found to lower cerebral Aβ levels and inhibit inflammatory activation in the brain, highlighting the gut-brain axis’s role in AD pathophysiology^68^.

#### Innate Immune Signaling Pathways

The NOD-like receptor signaling pathway, particularly the NLRP3 inflammasome, promotes the release of proinflammatory cytokines such as IL-1β and IL-18, exacerbating neuroinflammation and contributing to AD progression^69^. This pathway is further activated by dysregulated ions like K+ and Ca2+, prevalent in AD, which heightens the inflammatory response^69^. TLR signaling, notably through TLR3 and TLR4, influences Aβ dynamics and mediates neuroinflammation. TLR4 activation by Aβ in microglia triggers proinflammatory cytokine production, leading to amyloid-dependent neuronal death^70^. Similarly, dysregulated TLR pathways activate NF-κB and MAPK pathways, resulting in further inflammation and apoptosis^71^. The RLR pathway, although primarily known for antiviral responses, may exacerbate AD by promoting chronic inflammation through cytokine production. Lastly, T cells, particularly CD8+ T cells, infiltrate the brain and cerebrospinal fluid of AD patients, displaying increased expression of inflammatory pathways and significant clonal expansion^72^. Studies have demonstrated that T cells, especially cytotoxic T cells, are markedly increased in areas with tau pathology, correlating with neuronal loss and dynamically transforming from activated to exhausted states, which indicates their involvement in neurodegeneration^73^. Furthermore, the depletion of T cells has been shown to block tau-mediated neurodegeneration, suggesting that T cell activity directly contributes to disease progression^73^.

C-type lectin receptors (CLRs), such as CLEC-2, are found on the surface of platelets and are involved in the regulation of intestinal barrier function through their interaction with zonulin, a key modulator of intestinal permeability. Elevated levels of CLEC-2 and zonulin have been observed in patients with mild cognitive impairment and AD, suggesting a link between gut permeability and AD pathology. These elevated levels are also associated with reduced cognitive function as measured by the Mini-Mental State Examination score^74^. Additionally, CLRs are expressed by myeloid cells and recognize pathogen-associated molecular patterns and damage-associated molecular patterns, initiating immune responses that can contribute to inflammation, a known risk factor for AD^75^. Specifically, the scavenger receptor with C-type lectin (SRCL) has been implicated in the clearance of Aβ. SRCL is upregulated in astrocytes and vascular cells in AD patients and mouse models, suggesting its role in binding and clearing Aβ, thereby potentially mitigating AD progression^76^. Furthermore, the Dectin-1 cluster of CLRs, which includes receptors like CLEC-2, is involved in various pathophysiological processes, including inflammation and immune regulation, both of which are critical in the context of AD.

#### Cellular Processes

Neuroactive ligand-receptor interactions maintain brain homeostasis and regulate neurotransmitter systems, inflammatory responses, and neuroprotective mechanisms. In AD, disruptions in ligand-receptor networks, particularly those involving inflammatory pathways, are significant. For instance, microglial receptors interact with danger-associated molecular patterns to clear neurotoxic substances like Aβ and hyperphosphorylated tau^77,78^. Impairments in these interactions reduce the clearance of these toxic proteins, exacerbating disease progression. Chemokines and their receptors, part of neuroactive ligand-receptor interactions, have a dual role in AD: promoting neuroprotection and synaptic plasticity under normal conditions but leading to chronic inflammation when overexpressed, further contributing to Aβ aggregation and tau hyperphosphorylation^79^. Neurotransmitter receptors, such as cholinergic, glutamatergic, and serotonergic receptors, are also modulated in response to AD pathology, affecting cognitive functions and contributing to symptoms like memory loss and cognitive decline^80^. Multi-target-directed ligands that modulate these neurotransmitter systems show promise in providing symptomatic relief and potentially modifying disease progression by targeting multiple pathways involved in AD, including neuroinflammation and oxidative stress^81^. Therefore, understanding and targeting neuroactive ligand-receptor interactions offer significant potential for developing therapeutic strategies to combat AD.

TRP channels, such as TRPV1 and TRPA1, are implicated in the regulation of inflammatory processes in the brain. TRPV1, a non-selective cation channel, is involved in neuroinflammation and has been shown to influence microglial function. Activation of TRPV1 can rescue microglial dysfunction, restore metabolic impairments, and enhance immune responses, thereby reducing amyloid pathology and memory deficits in AD models^82,83^. On the other hand, TRPA1 channels, predominantly expressed in astrocytes, are activated by Aβ and mediate Ca2+ influx, which in turn triggers the production of pro-inflammatory cytokines and activation of transcription factors such as NF-κB and NFAT. Inhibition of TRPA1 channels reduces Aβ-induced inflammation and behavioral dysfunction, highlighting their role in AD pathogenesis^84^. Additionally, TRPC channels, particularly TRPC6, have been implicated in AD development, suggesting that TRP channels broadly contribute to the disease through their involvement in calcium homeostasis and glial cell activation^85^. The regulation of these channels by inflammatory mediators underscores their potential as therapeutic targets. Pharmacological modulation of TRP channels, such as using TRPV1 agonists, has shown promise in alleviating AD symptoms by reducing neuroinflammation and improving cellular functions^82,83^.

#### Bacterial Infections

Studies show tuberculosis (TB) patients have a higher risk of AD, potentially due to TB-induced chronic inflammation, which is a known AD risk factor^86^. Treatments for TB, like the BCG vaccine and rifampicin, show potential in modulating immune responses and reducing neuroinflammation, which could slow AD progression^87^. Again, maintaining gut and managing bacterial infections promptly, might mitigate risks for AD.

#### Viral Infections

Measles virus (MeV) has been implicated in neurodegenerative processes, as seen in subacute sclerosing panencephalitis, where persistent infection leads to neurofibrillary tangle formation, similar to AD^88^. This suggests that viral infections can contribute to neurodegenerative changes. The immune response to MeV, characterized by prolonged virus clearance and immunosuppression, may create a chronic inflammatory environment conducive to neurodegeneration^89^. Liver function markers, such as elevated AST to ALT ratios, are associated with AD diagnosis and cognitive dysfunction, suggesting metabolic disturbances linked to liver function influence AD pathophysiology^90^.

Hepatitis B virus (HBV), though less directly linked to AD, has been explored for its potential therapeutic avenues. Innovative research using HBV core protein to develop a vaccine targeting truncated tau proteins showed promising results in reducing tau pathology and cognitive deficits in a mouse model^91^.

#### Parasitic Infections

Chagas disease is primarily known for its impact on cardiac and gastrointestinal systems, but its potential role in the pathogenesis of AD can be inferred through its immunological and inflammatory mechanisms. The chronic infection with T. cruzi leads to a persistent inflammatory response and structural derangement in cardiac tissues, which is a hallmark of Chagas heart disease^92^. This inflammatory process is driven by the host’s immune response to the parasite, involving altered immunoregulatory mechanisms and pathogen persistence. Similarly, AD is characterized by neuroinflammation, where the accumulation of misfolded proteins activates an innate immune response, releasing inflammatory mediators that exacerbate the disease^93^. The cGAS–STING signaling pathway, which triggers type-I interferon-mediated neuroinflammation, is a critical component in AD pathogenesis^93^. Given that Chagas disease involves significant immune activation and inflammation, it is plausible that T. cruzi infection could influence neuroinflammatory pathways similar to those seen in AD. Additionally, the vascular pathogenesis of Chagas disease, involving functional changes in vasoactive peptides like endothelin-1 and kinins, could further contribute to neurovascular dysfunction, a known factor in AD progression^94^. Thus, while direct evidence linking Chagas disease to Alzheimer’s disease is limited, the shared mechanisms of chronic inflammation and immune dysregulation provide a basis for further investigation into their potential connection.

## 4. Discussion and conclusion

This study introduces mosGraphGPT, a generative pre-trained model designed for the integration and interpretation of multi-omics data. The primary aim was to enhance the understanding of Alzheimer’s disease (AD) pathogenesis by identifying significant signaling pathways and potential biomarkers through advanced graph neural networks (GNNs). The novel approach leverages extensive pre-training capabilities to capture complex gene-gene and gene-cell interactions with high accuracy and contextual relevance.

The integration of multi-omics data is crucial for understanding the intricate and multi-layered biological processes underlying complex diseases such as AD. Traditional models often struggle with the variability in gene expression and the diverse conditions of cell types. In contrast, foundation models like mosGraphGPT learn generalized representations from large-scale datasets, capturing complex interactions that simpler models cannot. This study utilized multi-omics datasets from UCSC Xena, encompassing epigenomics, genomics, transcriptomics, and proteomics data. The comprehensive dataset included 3592 cancer patients, 2121 genes, 8484 node entities, 19751 protein-protein interactions, and 26114 relations.

The experimental evaluation demonstrated that mosGraphGPT significantly improved disease classification accuracy and interpretability by uncovering disease targets and signaling interactions. The model achieved an average prediction accuracy of 75.09% on the ROSMAP AD dataset, outperforming other graph neural networks such as GCN, GAT, GIN, and UniMP. The results indicate the feasibility of patient outcome prediction using a GNN with a small set of core signaling pathways genes. The model’s ability to identify biomarkers and key signaling pathways was validated through attention mechanisms and statistical analyses. The integration of multi-omics data allowed for the identification of molecular mechanisms involving crucial molecular targets and signaling pathways. Notably, pathways such as the p53 signaling pathway, NF-kappa B signaling, and MAPK signaling were highlighted as significant in the context of AD. These findings are consistent with existing literature, underscoring the importance of these pathways in neurodegenerative diseases.

The findings from this study have significant implications for the field of bioinformatics and precision medicine. The ability of mosGraphGPT to integrate and interpret multi-omics data at a granular level provides a robust framework for understanding complex diseases. Future research could expand this model to other diseases and incorporate additional omics data to further refine the understanding of disease mechanisms and therapeutic targets. Additionally, the application of such models in clinical settings could enhance the precision of diagnostic and therapeutic strategies, paving the way for more personalized medicine approaches.

In conclusion, mosGraphGPT represents a significant advancement in the integration and interpretation of multi-omics data. By leveraging the extensive capabilities of generative pre-trained models and graph neural networks, this study has provided valuable insights into the molecular mechanisms of Alzheimer’s disease. The findings highlight the potential of such models to revolutionize the field of bioinformatics and precision medicine, offering a powerful tool for the study of complex diseases.

## Acknowledgement

This study was partially supported by NIA R56AG065352, 1R21AG078799-01A1, 1RM1NS132962-01, 1R01LM013902-01A1.

## Appendix

### Section A

**Figure S1.**
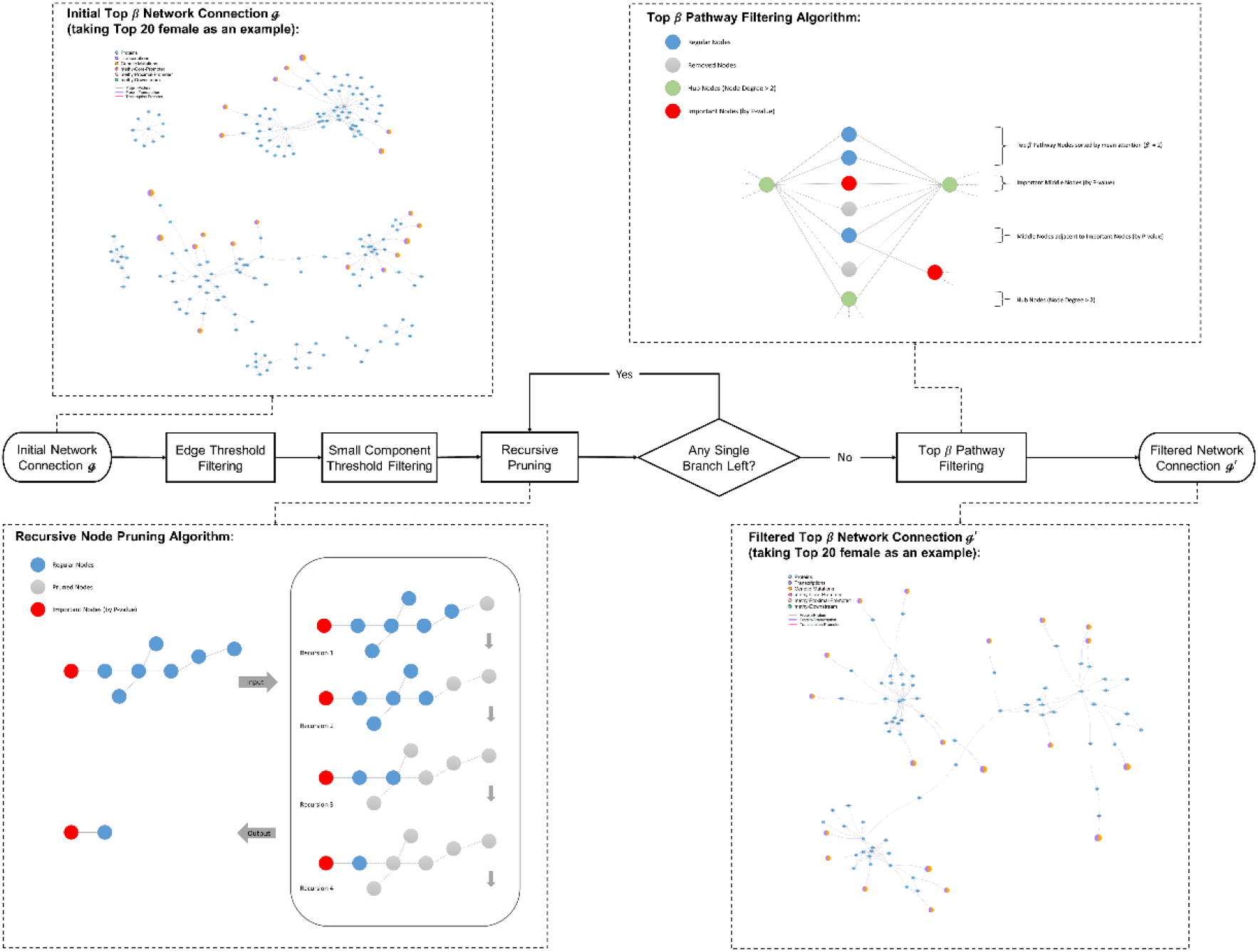
Diagram of the pruning procedures for core signaling networks visualization.

This pruning elucidates the systematic process of filtering and pruning network connections, essential for refining the network to its most significant components. The process begins with edge threshold filtering and small component threshold filtering. Edge threshold filtering eliminates edges that do not meet a specified significance level, thereby reducing the overall number of connections in the network. Small component threshold filtering removes smaller, less significant components from the network, focusing on the more substantial and potentially meaningful parts. Once the initial filtering is completed, a recursive node pruning algorithm is applied. This algorithm iteratively removes nodes that are deemed insignificant based on specific criteria, such as low connectivity or minimal contribution to the network’s overall structure. The purpose of this pruning is to simplify the network by eliminating nodes that do not add substantial value to the analysis, ensuring that only the most relevant nodes are retained.

After pruning, the algorithm checks whether any single branches remain within the network. A single branch is defined as a linear path with no bifurcations, which might not be as informative in the context of complex network structures. If a single branch is detected, pathway filtering is performed. Pathway filtering ensures that the remaining connections form biologically relevant pathways, thereby enhancing the network’s interpretability and utility for further analysis. The final result is a filtered network connection that retains the most significant elements, providing a clearer and more focused representation of the network’s structure. The diagram also includes examples of the initial and filtered network connections, illustrating the transformation that occurs through each stage of the process. Detailed steps of the recursive node pruning and pathway filtering algorithms are provided, enhancing the understanding of the methodology employed.

This meticulous approach to filtering and pruning ensures that the resulting network is not only simpler but also more biologically meaningful. By focusing on the most significant connections and pathways, researchers can achieve more accurate and insightful analyses, ultimately contributing to a deeper understanding of the underlying biological processes.

